# VDAC1 regulates stress-associated matrix localization of DJ-1 to support mitochondrial homeostasis and neuronal survival

**DOI:** 10.64898/2026.08.27.747597

**Authors:** Jessica Taday, Doo Soon Im, Sarah J. Hewitt, Gaurav Kaushik, Steve M. Callaghan, Ujval Anilkumar, Marisa Brini, Daniel Figeys, Zafran Khan, Nargis Khan, Ruth S. Slack, David S. Park, Alvin Joselin

**Affiliations:** Department of Clinical Neurosciences, Hotchkiss Brain Institute, Cumming School of Medicine, University of Calgary, Calgary, AB T2N 4N1, Canada; Department of Cellular and Molecular Medicine, Faculty of Medicine, Brain and Mind Research Institute, University of Ottawa, Ottawa, K1H 8M5, Canada; Department of Pharmaceutical and Pharmacological Sciences, University of Padova, Padova, 35131, Italy; Ottawa Institute of Systems Biology (OISB), University of Ottawa, Ottawa, K1H 8M5, Canada; Quadram Institute Bioscience, Norwich Research Park, and University of East Anglia, Norwich, Norfolk, UK; Department of Microbiology, Immunology and Infectious Diseases, Snyder Institute for Chronic Diseases Cumming School of Medicine, University of Calgary, Calgary, AB T2N 4N1, Canada

**Keywords:** Parkinson’s disease, VDAC1, DJ-1, mitochondria, bioenergetics

## Abstract

DJ-1 is a redox-sensitive protein implicated in early-onset Parkinson’s disease, and its mitochondrial localization protects against oxidative stress, but the mechanisms regulating its submitochondrial targeting and functional impact on mitochondrial integrity remain poorly understood. We identify voltage-dependent anion channel 1 (VDAC1) as a regulator of the submitochondrial distribution of DJ-1 during stress. Endogenous DJ-1 interacted with VDAC1, and loss of VDAC1 reduced stress-induced DJ-1 accumulation within the mitochondrial matrix. VDAC1-deficient neurons exhibited mitochondrial fragmentation, impaired oxidative phosphorylation, reduced ATP levels, altered reactive oxygen species (ROS) responses, and increased sensitivity to MPP⁺. Matrix-targeted, but not outer-membrane-targeted, DJ-1 rescued basal, ATP-linked, and maximal respiration, improved mitochondrial morphology, and enhanced neuronal survival. ATP synthase inhibition also rapidly increased mitochondrial DJ-1, suggesting bioenergetic stress promotes its mitochondrial accumulation. Our findings identify compartment-specific localization as a key determinant of DJ-1 function and establish VDAC1-dependent matrix targeting as a critical mechanism supporting mitochondrial integrity during stress.

## Introduction

Mitochondrial homeostasis is essential for cellular viability, particularly in metabolically active cells such as neurons. Disruptions in mitochondrial function underlie numerous neurodegenerative disorders, including Parkinson’s disease (PD), which is characterized by the progressive degeneration of dopaminergic (DA) neurons in the substantia nigra pars compacta (SNc). Although the majority of PD cases are sporadic, approximately 10% are associated with monogenic or familial forms of disease ^1^. Among the genes implicated in familial PD, DJ-1 (PARK7) is linked to early-onset disease and encodes a multifunctional protein involved in oxidative stress regulation, transcriptional control, and protein chaperoning ^2–4^.

Loss of DJ-1 function leads to mitochondrial fragmentation, increased reactive oxygen species (ROS), and heightened neuronal vulnerability to stress ^5,6^. Notably, the protective capacity of DJ-1 appears to depend, at least in part, on its ability to localize to mitochondria. Several studies have shown that DJ-1 accumulates at mitochondria in response to oxidative or metabolic stress; however, the mechanisms regulating this localization remain poorly defined ^2,7,8^. DJ-1 has also been reported to localize to different submitochondrial compartments, including the outer membrane (OMM), inner membrane (IMM), or mitochondrial matrix, suggesting compartment-specific functions ^2,9,10^.

The Voltage-Dependent Anion Channel 1 (VDAC1) is a central component of the outer mitochondrial membrane (OMM), where it facilitates the exchange of ions, nucleotides, and metabolites between the cytosol and the mitochondrial interior ^11–13^. While traditionally viewed as a metabolic gatekeeper, VDAC1 also functions as a regulatory hub by interacting with apoptotic and metabolic proteins such as Bcl-XL and hexokinase, thereby modulating mitochondrial permeability and cell death pathways ^14–16^. These dual roles position VDAC1 as a key mediator of mitochondrial health and stress responses. In a previous proteomic screen, we identified DJ-1, a redox-sensitive protein with neuroprotective functions, as a putative VDAC1-interacting partner. This observation led us to hypothesize that VDAC1 may influence DJ-1 submitochondrial localization and thereby affect mitochondrial integrity and cellular viability under stress.

In this study, we show that VDAC1 supports the stress-associated localization of DJ-1 to a protected mitochondrial compartment consistent with the matrix. Loss of VDAC1 reduces the stress-associated matrix localized pool of DJ-1 and is associated with mitochondrial dysfunction, including impaired oxidative phosphorylation, reduced ATP production, altered ROS responses, and mitochondrial fragmentation. Importantly, targeting DJ-1 specifically to the matrix restores basal, ATP-linked, and maximal mitochondrial respiration, improves mitochondrial morphology and enhances neuronal survival in VDAC1-deficient neurons. These findings define a previously unrecognized VDAC1-dependent axis that links mitochondrial stress, compartment-specific DJ-1 localization, and neuronal mitochondrial bioenergetics and integrity. This mechanism may be relevant to neuronal vulnerability in PD and other contexts of mitochondrial dysfunction.

## Results

### VDAC1 deficiency disrupts stress-induced mitochondrial accumulation of DJ-1

We previously employed a systems biology approach using an unbiased mass spectrometry screen to generate a large-scale human protein–protein interaction map ^17^. As part of this analysis, several candidate DJ-1-interacting proteins were identified. To validate these interactions, we performed additional biochemical experiments ^18^. Among the candidate interactors identified was VDAC1, a mitochondrial outer membrane protein. To validate this association, we first performed pull down assays in HEK293 cells transfected with GST or GST-DJ-1 constructs. Cell lysates were incubated with glutathione-Sepharose beads, and Western blot analysis revealed that VDAC1 was pulled down specifically in the presence of GST-DJ-1 but not GST alone, supporting a selective association between DJ-1 and VDAC1 (Figure 1A).

**Figure 1.**
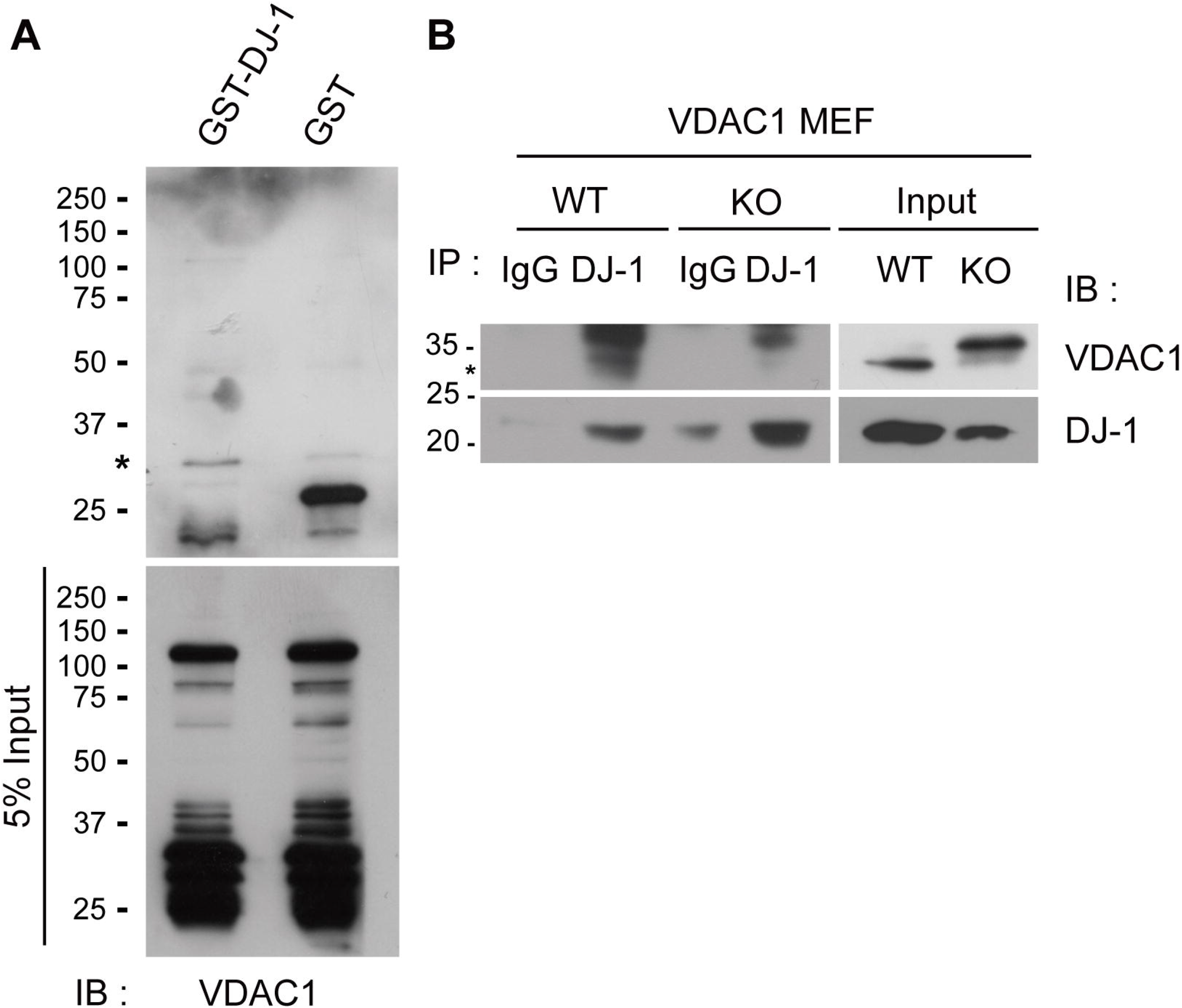
Biochemical validation of the association between DJ-1 and VDAC1. (A) GST pull-down demonstrating the association of GST-DJ-1 with endogenous VDAC1 in HEK293 cells. HEK293 cells were transfected with GST or GST-DJ-1 constructs, and lysates were subjected to GST pull-down using glutathione Sepharose beads followed by immunoblotting for VDAC1. The asterisk indicates the VDAC1-specific band. Lower panel shows input lysates immunoblotted for VDAC1. Representative immunoblots from n = 2 independent experiments. (B) Endogenous association of DJ-1 and VDAC1 in mitochondrial-enriched fractions from VDAC1 WT and KO MEFs. Mitochondrial-enriched lysates were immunoprecipitated with anti-DJ-1 antibody or control IgG and immunoblotted for VDAC1 and DJ-1. Input lysates are shown for VDAC1 and DJ-1. Representative immunoblots from n = 3 independent experiments.

To further confirm this interaction in a mitochondrial context, we isolated mitochondrial-enriched fractions from VDAC1 wild-type (WT) and knockout (KO) murine embryonic fibroblasts (MEFs). IP was performed using an anti–DJ-1 antibody or IgG control, followed by probing for VDAC1. W e observed that VDAC1 co-precipitated with DJ-1 in WT MEFs but not in KO cells, supporting a specific endogenous association between DJ-1 and VDAC1 (Figure 1B). We also detected DJ-1 in VDAC1 immunoprecipitates from embryonic brain lysate, but not in IgG control samples (Supplementary Figure 1). Although this experiment was performed once, it is consistent with the association observed in HEK293 cells and mitochondrial-enriched MEF fractions. These findings are consistent with a previous report that also identified VDAC1 as a binding partner of DJ-1 ^19^.

### VDAC1 KO neurons have higher basal mitochondrial DJ-1 than WT neurons

We previously reported that DJ-1 accumulates in mitochondrial fractions in response to oxidative stress in MEFs and primary cortical neurons ^7^. Given the localization of VDAC1 to the outer mitochondrial membrane and its association with DJ-1, we next examined whether VDAC1 influences the regulated stress-induced mitochondrial accumulation of DJ-1. VDAC1 WT and KO MEFs were treated with 100 μM H₂O₂ for up to 3 h, followed by isolation of mitochondrial-enriched fractions. Western blotting showed a time-dependent increase in mitochondrial DJ-1 in WT MEFs following H₂O₂ exposure. In contrast, VDAC1 KO MEFs did not show a comparable stress-induced increase in mitochondrial DJ-1 (Figure 2A, B). These findings suggest that VDAC1 deficiency alters the regulated mitochondrial accumulation of DJ-1 in response to oxidative stress in MEFs.

**Figure 2.**
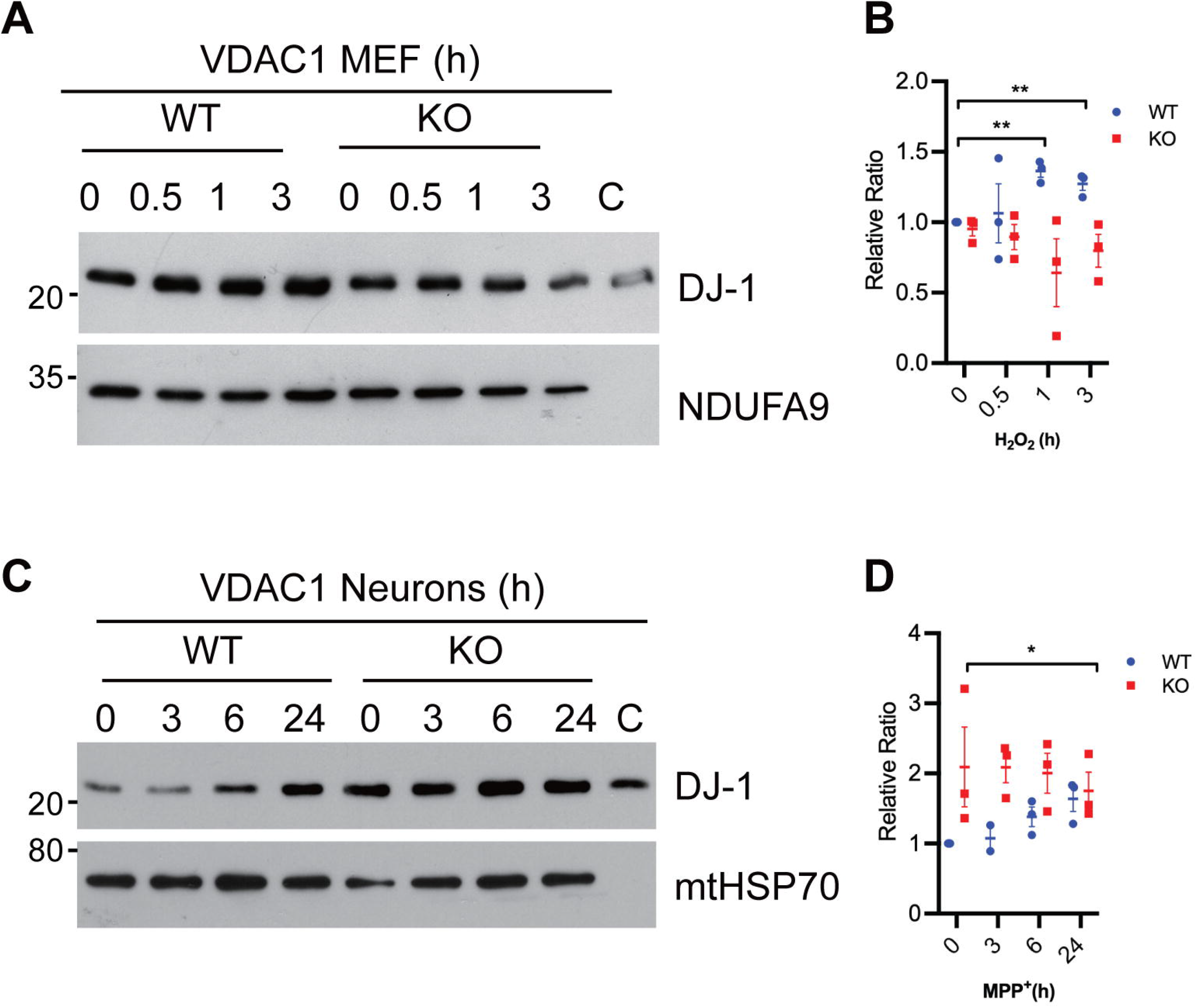
VDAC1 deficiency alters stress-induced mitochondrial accumulation of DJ-1. Western blot showing DJ-1 levels in mitochondrial-enriched fractions from VDAC1 WT and KO MEFs treated with 100 μM H₂O₂ for the indicated times (A) or primary cortical neurons treated with 10 μM MPP⁺ for the indicated times (C). NDUFA9 was used as a mitochondrial loading control for MEFs (A) and mtHSP70 for cortical neurons (C). Representative immunoblots from n = 3 independent experiments. (B) Quantification of mitochondrial DJ-1 levels shown in (A),. Data represent mean ± SEM (n = 3 independent experiments). Two-way ANOVA with Tukey’s multiple-comparisons test. **p < 0.01. (B and D) Quantification of mitochondrial DJ-1 levels shown in (A and C), normalized to NDUFA9 (B) and mtHSP70 (D). Data represent mean ± SEM (n = 3 independent embryo-derived cultures). WT and KO groups were compared at individual time points using two-tailed unpaired Student’s t-tests. p < 0.05.

We extended this analysis to primary cortical neurons derived from E14.5–15.5 VDAC1 WT and KO embryos. At DIV6, neurons were treated with 10 μM MPP⁺ (1-methyl-4-phenylpyridinium), the active toxic metabolite of MPTP commonly used to model Parkinsonian mitochondrial stress, for up to 24 hours in antioxidant-free media. Mitochondrial-enriched fractions were then isolated and analyzed by Western blot using antibodies against DJ-1 and the mitochondrial matrix protein mtHSP70. In WT neurons, mitochondrial DJ-1 increased significantly between 6 and 24 hours after MPP⁺ exposure. In contrast, VDAC1 KO neurons displayed elevated basal DJ-1 but did not exhibit a further increase after MPP⁺ treatment (Figure 2C, D). Together, these findings indicate that VDAC1 deficiency alters the regulated stress-associated mitochondrial accumulation of DJ-1 in both MEFs and primary cortical neurons. Notably, VDAC1 KO neurons exhibited elevated basal mitochondrial DJ-1 despite failing to show a further stress-induced increase.

### VDAC1 deficiency impairs neuronal bioenergetics

Considering that VDAC1 KO neurons exhibit elevated basal mitochondrial DJ-1, we hypothesized that this may reflect an underlying state of mitochondrial dysfunction. While VDAC1’s role in regulating mitochondrial metabolism has been studied in tissues such as liver and heart ^20–23^, its contribution to neuronal mitochondrial health remains unclear. Given the unique energetic demands of neurons, we sought to assess how VDAC1 deficiency affects neuronal mitochondrial function and stress responses.

To evaluate mitochondrial respiration, we cultured cortical neurons from E14.5–15.5 VDAC1 WT and KO embryos and measured oxygen consumption rate (OCR) using a Seahorse XF-24 analyzer. Cortical neurons derived from three independent embryos from the same litter were analyzed at baseline and following sequential addition of oligomycin (ATP synthase inhibitor), FCCP (uncoupler), and rotenone/antimycin A (complex I and III inhibitors). VDAC1 KO neurons exhibited a markedly reduced OCR compared to WT controls across all phases of the assay (Figure 3A). Quantitative analysis showed significant decreases in basal respiration, ATP-linked respiration, maximal respiration, and spare respiratory capacity in KO neurons (Figure 3B–E), indicating a broad impairment in oxidative phosphorylation.

**Figure 3.**
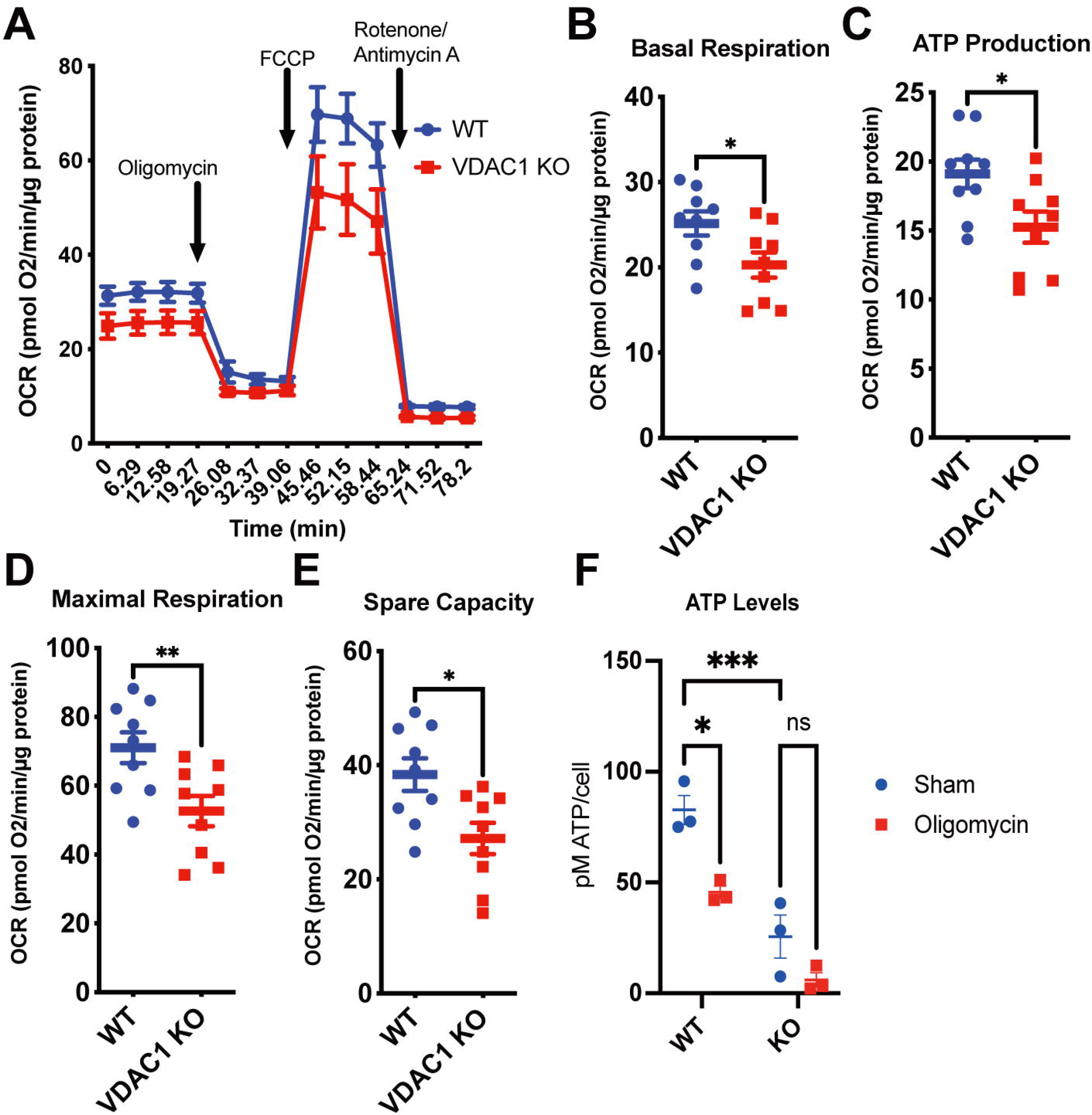
VDAC1 deficiency impairs mitochondrial respiration and ATP production in primary cortical neurons. (A) Representative oxygen consumption rate (OCR) traces from VDAC1 WT and KO primary cortical neurons (DIV9–10) analyzed using the Seahorse XF24 Analyzer. Sequential injections of oligomycin (500 ng/mL), FCCP (2 μM), and rotenone (0.5 μM)/antimycin A (1 μM) are indicated. Representative traces from at least three independent embryo-derived cultures. (B–E) Quantification of basal respiration (B), ATP-linked respiration (C), maximal respiration (D), and spare respiratory capacity (E) from the Seahorse assay shown in (A). Data represent mean ± SEM (n = 3 biological replicates, each representing an independent embryo-derived culture). Two-tailed Student’s *t* test. *p < 0.05; **p < 0.01. (F) Total ATP levels in VDAC1 WT and KO primary cortical neurons treated with or without 10 μM oligomycin. ATP levels were measured using a luciferase-based assay and normalized to cell number. Data represent mean ± SEM (n = 3 biological replicates). Two-way ANOVA with Tukey’s multiple-comparisons test. *p < 0.05; ***p < 0.001.

To determine whether this defect translated into altered ATP availability, we measured total ATP in WT and KO neurons using a luciferase-based assay. VDAC1 KO neurons had significantly lower basal ATP levels than WT neurons (Figure 3F). Oligomycin reduced ATP levels in WT neurons, whereas the reduction was substantially smaller in VDAC1 KO neurons. Collectively, these findings indicate that VDAC1 deficiency results in impaired mitochondrial oxidative phosphorylation in cortical neurons, leading to reduced ATP production and a basal bioenergetic deficit with limited additional sensitivity to acute ATP synthase inhibition.

### VDAC1 deficiency disrupts mitochondrial morphology and alters ROS responses in neurons

Given the reduction in mitochondrial respiration and ATP levels observed in VDAC1-deficient neurons, we next asked whether these functional changes were accompanied by alterations in mitochondrial morphology. Mitochondrial shape provides a useful readout of organellar state, and a shift toward shorter, more fragmented mitochondria is commonly associated with cellular stress and mitochondrial dysfunction ^24–26^.

To evaluate mitochondrial morphology, we stained DIV6 VDAC1 WT and KO cortical neurons with Tom20 and visualized mitochondria using confocal microscopy. Compared to the elongated, tubular mitochondrial networks observed in WT neurons, mitochondria in KO neurons appeared shorter and more fragmented (Figure 4A). To quantify this more rigorously, we analyzed mitochondrial length distributions. Mitochondria were binned into five length categories (<0.5 μm, 0.5–1 μm, 1.01–2 μm, 2.01–3 μm, and >3 μm), and the percentage of mitochondria falling into each category was calculated. VDAC1 KO neurons showed a significant increase in the proportion of short mitochondria (<1 μm), with a corresponding decrease in the population of mitochondria longer than 1 μm compared to WT neurons (Figure 4B). This shift toward shorter mitochondrial length is consistent with increased fragmentation in the absence of VDAC1.

**Figure 4.**
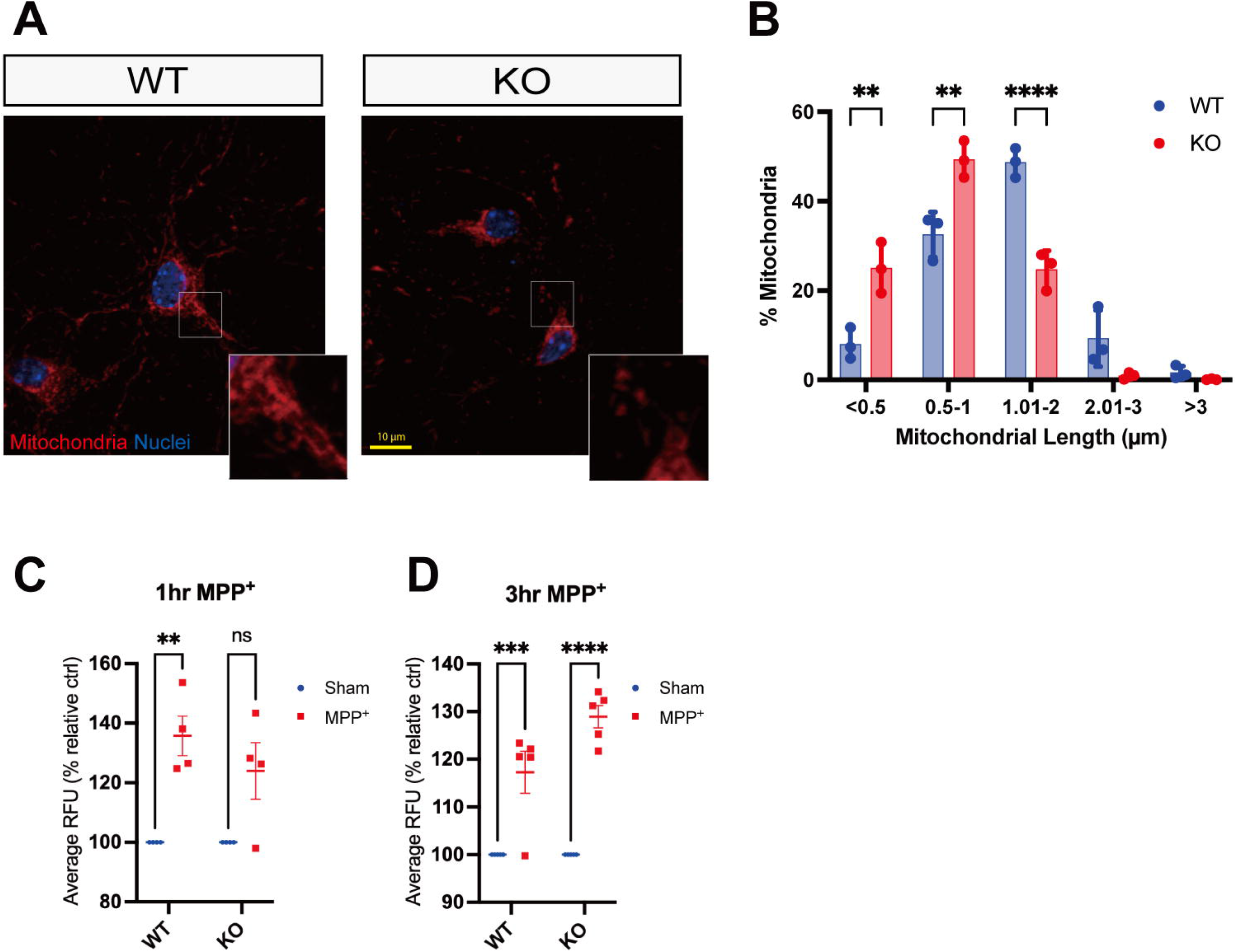
VDAC1 deficiency disrupts mitochondrial morphology and alters ROS responses. (A) Representative confocal images of VDAC1 WT and KO primary cortical neurons at DIV6 stained for Tom20 and Hoechst. Scale bar: 10 μm. (B) Quantification of mitochondrial length distribution in VDAC1 WT and KO neurons. Mitochondria were binned into five length categories: <0.5 μm, 0.5–1 μm, 1.01–2 μm, 2.01–3 μm, and >3 μm. Data represent mean ± SEM (n = 3 biological replicates; ≥500 total mitochondria per condition). Two-way ANOVA with Sidak’s post hoc test. **p < 0.01, ***p < 0.0001. (C–D) Intracellular ROS levels in VDAC1 WT and KO primary cortical neurons treated with 10 μM MPP⁺ for 1 h (C) or 3 h (D) in antioxidant-free medium. ROS levels were measured using CM-H₂DCFDA fluorescence. Quantification was performed from 4–6 randomly selected fields per condition. Data are mean ± SEM (n = 3–5 independent embryo-derived cultures). Two-way ANOVA with Bonferroni post hoc test. *p < 0.05, **p < 0.01, ***p < 0.001, ****p < 0.0001.

To assess redox homeostasis, we treated WT and KO neurons with 10 μM MPP⁺ for 1 or 3 hours and measured intracellular ROS using CM-H₂DCFDA. At 1 h (Figure 4C), KO neurons showed reduced ROS levels compared to WT, but at 3 h (Figure 4D), ROS levels were significantly higher, suggesting a dysregulated oxidative stress response. Taken together, these findings indicate that VDAC1 deficiency is associated with mitochondrial fragmentation and an altered ROS response to MPP⁺ in primary cortical neurons.

### VDAC1 regulates submitochondrial localization of DJ-1 during oxidative stress

Given the observed mitochondrial dysfunction in VDAC1-deficient neurons, we sought to investigate whether altered DJ-1 localization contributes to these defects. Although DJ-1 has been reported to localize to mitochondria in several studies ^2,7–9^, its mechanism of mitochondrial targeting remains unclear due to the absence of conventional mitochondrial localization signals. Furthermore, the precise submitochondrial compartment in which DJ-1 resides is debated, with reports describing its presence on the outer mitochondrial membrane (OMM), inner mitochondrial membrane (IMM), and/or matrix ^9,20–23^.

To clarify DJ-1’s submitochondrial localization and its dependence on VDAC1, we subjected mitochondrial-enriched fractions from VDAC1 WT and KO MEFs to proteolytic digestion with trypsin for up to 120 minutes under oxidative stress conditions (100 μM H₂O₂). In this assay, OMM proteins are rapidly digested, whereas matrix proteins remain protected unless the membrane integrity is compromised over time. In WT MEFs, DJ-1 was largely resistant to trypsin digestion in both control and H₂O₂-treated conditions, with protection kinetics similar to mtHSP70, consistent with localization within a protected mitochondrial compartment, likely the matrix (Figure 5A–B). In contrast, while DJ-1 in VDAC1 KO MEFs remained protected under basal conditions (Figure 5C), it became increasingly susceptible to proteolysis under oxidative stress (Figure 5D). These findings suggest that VDAC1 deficiency compromises stress-associated localization of DJ-1 to a protected mitochondrial compartment consistent with the matrix. As a control for protease accessibility, mitochondrial fractions were treated with Triton X-100 prior to trypsin digestion, which rendered DJ-1, mtHSP70, and other mitochondrial markers susceptible to proteolysis (Supplementary Figure 2A–B).

**Figure 5.**
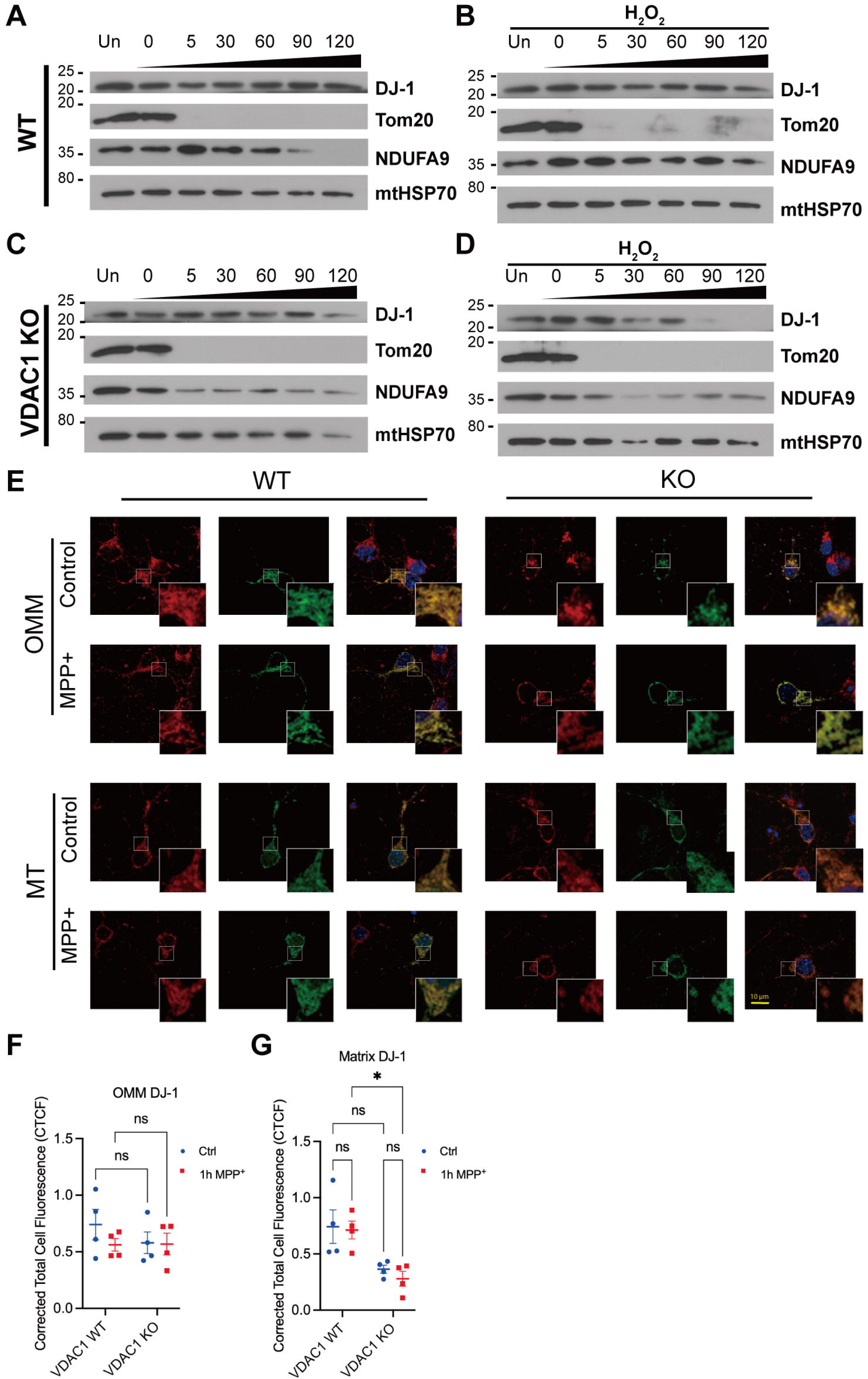
VDAC1 deficiency alters stress-associated matrix localization of DJ-1. (A–D) Trypsin digestion of mitochondrial-enriched fractions isolated from VDAC1 WT and KO MEFs treated with or without 100 μM H₂O₂ for 3 h. Fractions were digested with 25 μg/mL trypsin for 0–120 min and analyzed by Western blotting for DJ-1, Tom20, NDUFA9, and mtHSP70. Representative immunoblots from n = 3 independent experiments. (E) GFP complementation assay in VDAC1 WT and KO cortical neurons. Neurons were co- transfected with DJ-1-S11 and either OMM-targeted GFP1–10 or matrix-targeted GFP1–10 constructs, followed by treatment with or without 10 μM MPP⁺ for 3 h. Cells were stained for Tom20 and Hoechst. Insets show enlarged regions for all panels. Scale bar: 10 μm. Representative of n = 4 independent experiments. (F–G) Quantification of Corrected Total Cell Fluorescence (CTCF) from OMM-targeted GFP complementation (F) and matrix-targeted GFP complementation (G). Data represent mean ± SEM (n = 4 biological replicates, averaging 5-8 cells per condition). Two-way ANOVA with Tukey’s post hoc test. *p < 0.05.

To further probe this question in neurons, we employed a GFP complementation assay, which allows for precise localization of DJ-1 to specific mitochondrial sub-compartments ^9^. In this system, the green fluorescent protein (GFP) is split into two fragments: GFP1-10 and the S11 β-strand. GFP1-10 is targeted to either the OMM or mitochondrial matrix, while DJ-1 is fused to S11. When DJ-1-S11 reaches the same compartment as the targeted GFP1-10, the fragments reassemble and fluoresce, allowing us to infer DJ-1 localization based on compartment-specific GFP complementation. VDAC1 WT and KO cortical neurons were transfected with either OMM-GFP1–10 or MT-GFP1–10 constructs, together with WT-DJ1-S11, and treated with 10 μM MPP⁺ for 3 hours.

Representative confocal images demonstrated robust reconstituted GFP fluorescence in both OMM- and matrix-targeted constructs in WT neurons under both control and stress conditions (Figure 5E). Quantification of Corrected Total Cell Fluorescence (CTCF) confirmed no significant differences in OMM-targeted DJ-1 fluorescence between groups (Figure 5F), indicating that VDAC1 is not required for OMM association of DJ-1. Matrix-localized DJ-1 signal was significantly reduced in VDAC1 KO neurons (n = 4), especially following MPP⁺, consistent with impaired stress-associated matrix localization (Figure 5G). These findings indicate that VDAC1 supports efficient stress-associated matrix localization of DJ-1 during MPP+-induced stress. These findings support a role for VDAC1 in directing DJ-1 to the matrix during stress.

### Bioenergetic stress promotes mitochondrial accumulation of DJ-1

We previously reported that DJ-1 localizes to mitochondria in response to oxidative stress in neurons ^7^, but the upstream signal contributing to this translocation remain unclear. To determine whether exogenous oxidative stress alone accounts for DJ-1 mitochondrial accumulation, we pretreated primary cortical neurons with the antioxidants N-acetylcysteine (NAC) for 2 hours prior to H₂O₂ stimulation. NAC treatment alone increased mitochondrial DJ-1 levels, with no further increase observed following H₂O₂ exposure (Figure 6A). These results suggest that DJ-1 mitochondrial accumulation is not simply proportional to exogenous oxidative stress and may also be influenced by changes in cellular redox or metabolic state.

**Figure 6.**
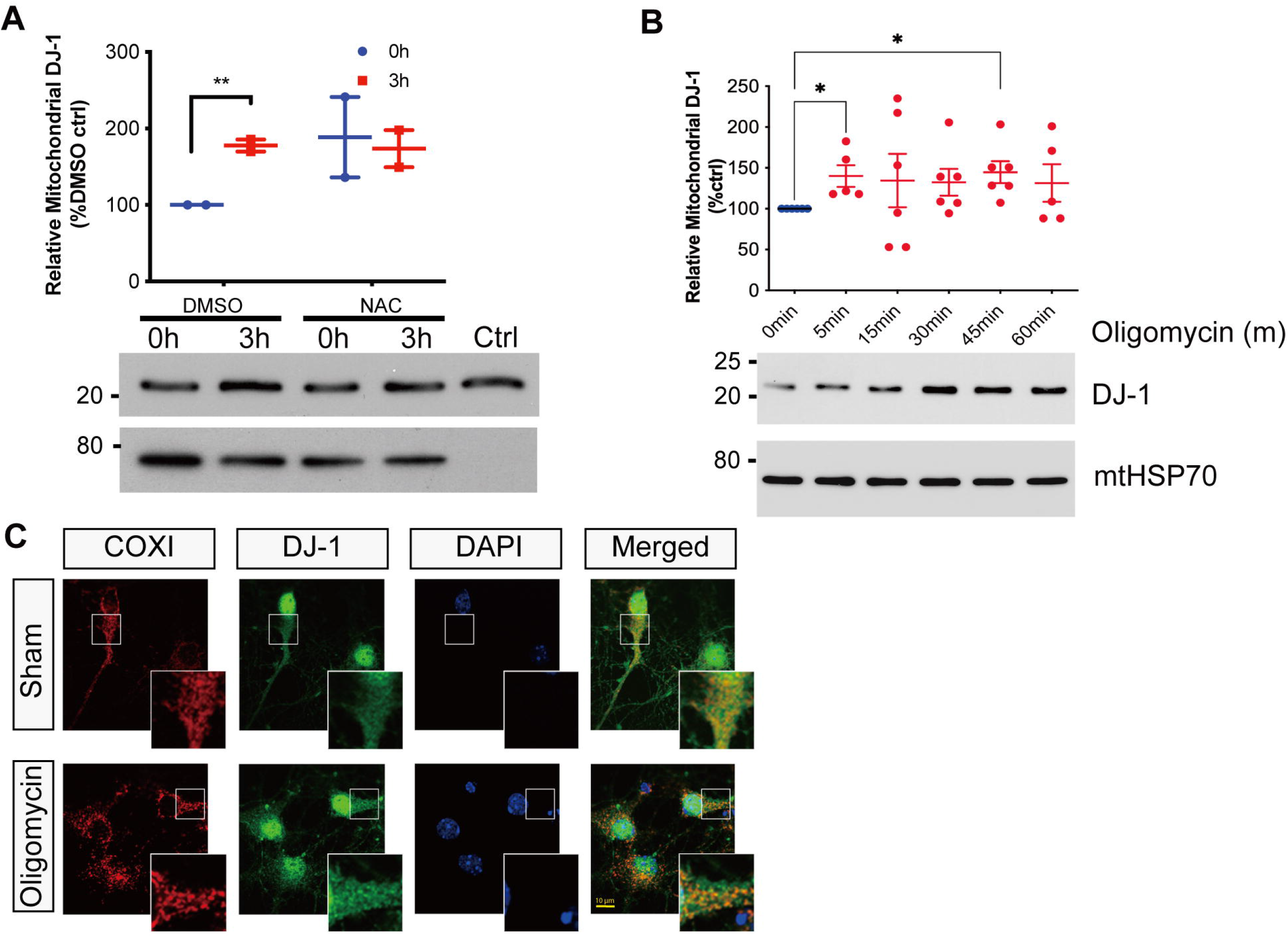
Bioenergetic stress promotes mitochondrial accumulation of DJ-1. (A) CD-1 primary cortical neurons were pretreated with 2 mM N-acetylcysteine (NAC) for 2 h, followed by treatment with 30 μM H₂O₂ for the indicated times. Mitochondrial-enriched fractions were immunoblotted for DJ-1 and mtHSP70. Representative immunoblots from n = 3 independent experiments. (B) CD-1 primary cortical neurons were treated with 10 μM oligomycin for the indicated times in antioxidant-free medium. Mitochondrial-enriched fractions were immunoblotted for DJ-1 and mtHSP70. Quantification of mitochondrial DJ-1 levels normalized to mtHSP70 is shown. Data represent mean ± SEM (n = 5–6 independent experiments). Unpaired two-tailed Student’s *t* test versus 0 min. *p < 0.05, **p < 0.01. (C) Representative confocal images of CD-1 primary cortical neurons (DIV6) treated with 10 μM oligomycin for 15 min and immunostained for DJ-1, COXI, and Hoechst. Insets show enlarged regions for all panels. Scale bar, 10 μm. Representative images from n = 2 independent experiments.

To test whether bioenergetic stress is sufficient to promote DJ-1 mitochondrial accumulation, we treated WT cortical neurons with 10 μM oligomycin, an ATP synthase inhibitor, for up to 60 minutes in antioxidant-free media. Western blot analysis of mitochondrial-enriched fractions revealed a rapid and sustained increase in mitochondrial DJ-1 levels, evident as early as 5 minutes after treatment and persisting throughout the time course (Figure 6B). Immunofluorescence staining further supported these findings, showing increased colocalization of endogenous DJ-1 with the mitochondrial marker COXI following oligomycin treatment (Figure 6C). Together, these findings indicate that inhibition of ATP synthase is sufficient to promote mitochondrial accumulation of DJ-1 and may help explain the elevated basal mitochondrial DJ-1 observed in VDAC1-deficient neurons (Figure 2).

### Matrix-targeted DJ-1 rescues mitochondrial respiration in VDAC1- deficient neurons

Given that bioenergetic stress promotes mitochondrial accumulation of DJ-1, we next asked whether directing DJ-1 to the matrix could functionally rescue the respiratory deficit observed in VDAC1 KO neurons. VDAC1 KO cortical neurons were transduced with AAV2/9-CBA-EGFP, AAV2/9-CBA-OMM-DJ1-myc or AAV2/9-CBA-MT-DJ1-myc, and oxygen consumption rate (OCR) was measured using a Seahorse XF96 Extracellular Flux Analyzer. KO neurons expressing GFP or OMM-targeted DJ-1 showed overlapping OCR profiles with visibly lower basal and drug-responsive oxygen consumption throughout the assay, whereas KO neurons expressing matrix-targeted DJ-1 (MT-DJ1) displayed an upward shift in OCR across all phases of the assay relative to GFP- and OMM-DJ1-expressing neurons (Figure 7A).

**Figure 7.**
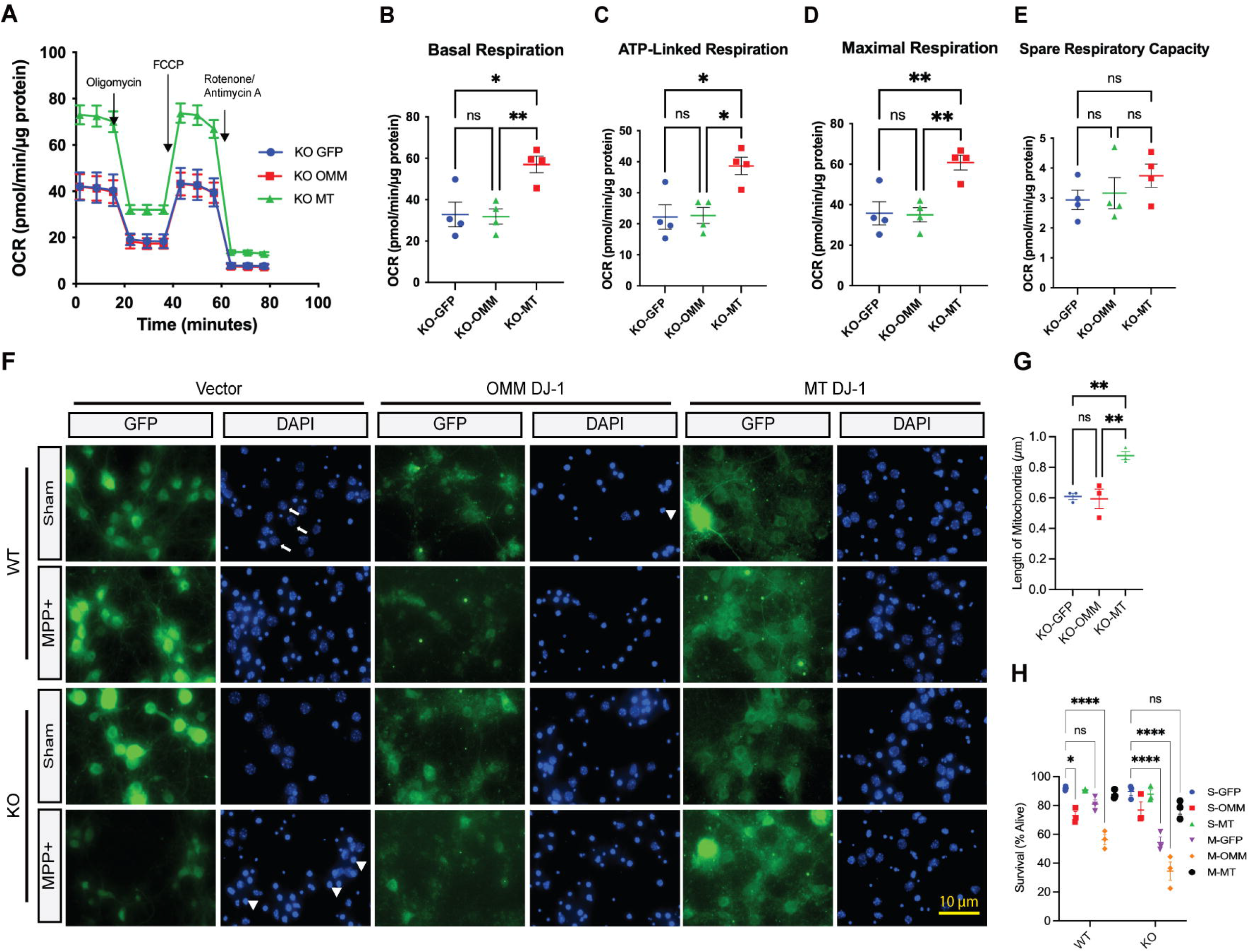
Matrix-targeted DJ-1 rescues mitochondrial respiration, improves mitochondrial morphology and enhances survival in VDAC1-deficient neurons. (A) Representative OCR traces from VDAC1 KO primary cortical neurons transduced with AAV2/9-CBA-EGFP, AAV2/9-CBA-OMM-DJ1-myc, or AAV2/9-CBA-MT-DJ1-myc and analyzed using the Seahorse XF96 Analyzer. Sequential injections of oligomycin (1 μM), FCCP (2 μM), and rotenone (0.5 μM)/antimycin A (0.5 μM) are indicated. Representative traces from n = 4 biological replicates. (B–E) Quantification of basal respiration (B), maximal respiration (C), ATP-linked respiration (D), and spare respiratory capacity (E) from the Seahorse assay shown in (A). Data represent mean ± SEM (n = 4 biological replicates). One-way ANOVA with Tukey’s multiple-comparisons test. *p < 0.05, **p < 0.01. (F) Representative images of VDAC1 WT and KO primary cortical neurons transduced with AAV2/3-CBA-GFP, AAV2/3-CBA-OMM-DJ1-GFP, or AAV2/3-CBA-MT-DJ1-GFP and treated with 10 μM MPP⁺ for 48 h. Neurons were stained with Hoechst to assess nuclear morphology. Arrows indicate live cells and arrowheads indicate dead cells. Scale bar, 10 μm. (G) Quantification of mitochondrial length in VDAC1 KO primary cortical neurons transduced with AAV2/3-CBA-GFP, AAV2/3-CBA-OMM-DJ1-GFP, or AAV2/3-CBA-MT-DJ1-GFP and treated with 10 μM MPP⁺. Data represent mean ± SEM (n = 3 biological replicates). One-way ANOVA with Tukey’s multiple-comparisons test. **p < 0.01. (H) Quantification of neuronal survival across genotype, treatment, and AAV transduction conditions. Data represent mean ± SEM (n = 3 independent embryo-derived cultures; ≥130 cells analyzed per condition). Two-way ANOVA with Tukey’s multiple- comparisons test. *p < 0.05, **p < 0.01, ***p < 0.001, ****p < 0.0001.

Quantitative analysis was consistent with this rescue. Basal respiration was significantly increased in MT-DJ1-expressing KO neurons compared with GFP controls, while OMM-DJ1 did not differ from GFP (Figure 7B). Similarly, maximal respiration and ATP-linked respiration were both significantly elevated in MT-DJ1-expressing neurons relative to GFP and OMM-DJ1 groups, with no significant difference between GFP and OMM-DJ1 (Figure 7C-D). In contrast, spare respiratory capacity remained low and did not differ significantly among GFP, OMM-DJ1, and MT-DJ1 groups (Figure 7E), Together, these findings indicate that matrix-targeted, but not OMM-targeted, DJ-1 improves multiple parameters of mitochondrial respiration in VDAC1-deficient neurons.

### Matrix-localized DJ-1 protects VDAC1-deficient neurons from MPP**⁺** -induced death

We next assessed whether DJ-1 submitochondrial targeting influences mitochondrial morphology and neuronal survival under stress. VDAC1 WT and KO cortical neurons were transduced with AAV2/3-CBA-GFP, AAV2/3-CBA-OMM-DJ1-GFP or AAV2/3-CBA-MT-DJ1-GFP. To first assess whether compartment-targeted DJ-1 could modify the mitochondrial morphology defect observed in VDAC1-deficient neurons, we measured mitochondrial length following MPP⁺ exposure. Expression of MT-DJ1 significantly increased average mitochondrial length in VDAC1 KO neurons compared with GFP controls, indicating reduced mitochondrial fragmentation, whereas OMM-DJ1 did not significantly improve mitochondrial length (Figure 7F).

We next assessed neuronal survival following treatment with 10 μM MPP⁺ for 48 hours. Following MPP⁺ treatment, VDAC1 KO neurons transduced with GFP showed reduced survival compared with sham-treated controls (Figure 7G–H). OMM-targeted DJ-1 did not rescue this loss of viability and was associated with a further reduction in survival following MPP⁺ exposure. In contrast, matrix-targeted DJ-1 significantly improved survival compared with GFP-transduced, MPP⁺-treated VDAC1 KO neurons. These findings suggest that DJ-1-mediated protection is associated with its submitochondrial targeting, with matrix-targeted DJ-1 conferring protection under MPP⁺ stress.

## Discussion

In this study, we identify VDAC1 as an important regulator of DJ-1 submitochondrial localization and mitochondrial stress adaptation. We show that DJ-1 associates with VDAC1, that loss of VDAC1 disrupts stress-associated accumulation of DJ-1 within a protected mitochondrial compartment consistent with the matrix, and that VDAC1-deficient neurons exhibit mitochondrial fragmentation, impaired respiration, reduced ATP production, and increased vulnerability to MPP⁺. Importantly, matrix-targeted DJ-1 improved mitochondrial respiration, morphology and neuronal survival in VDAC1-deficient neurons, supporting the interpretation that matrix-localized DJ-1 contributes to mitochondrial protection under stress.

Previous studies, including our own work, have shown that DJ-1 plays an important role in preserving mitochondrial health and that stress-induced localization of DJ-1 in the mitochondria promotes cell survival ^2,5,7,8,27^. VDAC1, often referred to as the primary gateway for mitochondrial metabolites to communicate with the rest of the cell, is also a key regulator of mitochondrial function and mitochondria-mediated apoptosis ^16^. Our findings connect these two mitochondrial stress pathways by suggesting that VDAC1 contributes to the compartment-specific localization of DJ-1 during mitochondrial stress. Although VDAC1 is localized to the outer mitochondrial membrane, its effects on DJ-1 localization are more likely to reflect broader roles in coordinating outer membrane permeability, contact-site organization, or bioenergetic signalling rather than direct import of DJ-1 into the matrix. The precise mechanism by which VDAC1 regulates this process remains to be determined.

Consistent with this possibility, our data further suggest that bioenergetic stress can promote mitochondrial accumulation of DJ-1. Oligomycin rapidly increased mitochondrial DJ-1 levels, indicating that ATP synthase inhibition is sufficient to promote this response. However, the relationship between ATP depletion, redox state, and DJ-1 localization is likely complex. NAC treatment did not block DJ-1 mitochondrial accumulation and instead increased mitochondrial DJ-1 levels on its own, suggesting that DJ-1 localization is not simply proportional to exogenous oxidative stress. These findings support a model in which DJ-1 responds to mitochondrial metabolic state, but they do not exclude contributions from redox-dependent signalling. This interpretation is consistent with the elevated basal mitochondrial DJ-1 observed in VDAC1-deficient neurons, while the MEF data indicate a loss of regulated stress-induced accumulation without a clear basal increase. Although the patterns differed between cell types, the common feature in both models was impaired stress-associated localization of DJ-1 to a protected mitochondrial compartment. The elevated basal mitochondrial DJ-1 observed in VDAC1-deficient neurons likely reflects differences in the basal bioenergetic state of neurons compared with MEFs rather than preserved stress-responsive trafficking.

The elevated basal mitochondrial DJ-1 observed in VDAC1-deficient neurons may therefore reflect a compensatory response to pre-existing mitochondrial stress rather than normal DJ-1 trafficking. This interpretation is consistent with the reduced respiration and ATP levels observed in VDAC1-deficient neurons, as well as the rapid mitochondrial accumulation of DJ-1 following oligomycin treatment. Importantly, the presence of DJ-1 in mitochondrial-enriched fractions does not necessarily indicate efficient matrix localization. Indeed, the trypsin-protection and split-GFP experiments suggest that VDAC1 deficiency compromises the stress-associated matrix-protected pool of DJ-1. Thus, VDAC1 loss may uncouple mitochondrial recruitment of DJ-1 from its productive compartment-specific localization within mitochondria.

Mitochondrial morphology is closely linked to mitochondrial function and cellular health, and disruptions in mitochondrial fusion, fission, and network organization have been associated with mitochondrial dysfunction and disease ^24,25,28^. In our study, VDAC1-deficient cortical neurons exhibited reduced oxygen consumption, decreased ATP levels, and shorter mitochondria compared with wild-type neurons. These findings indicate that loss of VDAC1 is associated with both bioenergetic impairment and mitochondrial fragmentation, consistent with a broader disruption of mitochondrial homeostasis. Given that mitochondrial dysfunction can increase ROS production, limit ATP availability, and compromise cell survival in neurodegenerative disease ^24,29,30^, these changes may contribute to the increased vulnerability of VDAC1-deficient neurons under stress.

Loss of DJ-1 has been reported to alter mitochondrial morphology and dynamics in MEF cells ^5^, resembling the mitochondrial fragmentation observed in VDAC1-deficient neurons in our study. These phenotypic similarities are consistent with the possibility VDAC1 and DJ-1 function within overlapping mitochondrial stress-response pathways. This idea is further supported by reports that DJ-1 can localize to multiple submitochondrial compartments under stress conditions, including the outer mitochondrial membrane, inner mitochondrial membrane, and matrix ^2,10,31,32^. Our findings help refine these observations by showing that DJ-1 localizes to a protected mitochondrial compartment consistent with the matrix in trypsin-protection and split-GFP complementation assays, and that this matrix-associated pool is reduced in VDAC1-deficient cells under stress. Notably, the split-GFP complementation assay did not show a generalized increase in mitochondrial DJ-1 signal under stress, despite previous reports from our group and others showing stress-induced mitochondrial accumulation of endogenous DJ-1. This discrepancy may reflect differences in cell type, stress paradigm, assay sensitivity, or the requirement for overexpressed DJ-1 fusion constructs in the split-GFP system. These observations suggest that stress-associated localization of DJ-1 may occur through multiple sequential or dynamic mitochondrial compartments rather than a single static localization state.

Consistent with this concept, restoring DJ-1 specifically to the mitochondrial matrix improved multiple aspects of mitochondrial function in VDAC1-deficient neurons. Matrix-targeted DJ-1 increased basal, ATP-linked, and maximal respiration, improved mitochondrial morphology, and enhanced neuronal survival following MPP⁺ exposure. In contrast, targeting DJ-1 to the outer mitochondrial membrane failed to rescue these phenotypes, highlighting that mitochondrial localization alone is insufficient and that compartment-specific localization is required for optimal function. Previous studies have reported stress-induced accumulation of DJ-1 at the outer mitochondrial membrane, whereas others have detected DJ-1 within internal mitochondrial compartments^2,8,10^. Our findings suggest that these observations are not necessarily contradictory. Although mitochondrial surface association may occur during oxidative stress, localization to a protected mitochondrial compartment consistent with the matrix appears to be required for optimal mitochondrial function and neuroprotection. Notably, matrix-targeted DJ-1 did not restore spare respiratory capacity, indicating that matrix-associated DJ1 contributes substantially to mitochondrial protection but does not fully account for the broader roles of VDAC1 in maintaining mitochondrial function. Furthermore, because matrix targeting was achieved using an engineered localization sequence, these experiments demonstrate the functional importance of matrix-associated DJ-1 but do not establish the mechanism by which endogenous DJ-1 reaches this compartment.

These findings raise several important questions for future study. Although our data support a role for VDAC1 in regulating the localization of DJ-1 to a protected mitochondrial compartment, the underlying mechanism remains to be defined. VDAC1 may influence DJ-1 localization through direct interaction, altered mitochondrial contact-site organization, changes in outer membrane permeability, or secondary effects on mitochondrial bioenergetics. Future studies will be required to determine how endogenous DJ-1 reaches this compartment and whether this pathway contributes to selective neuronal vulnerability in Parkinson’s Disease. In particular, studies using dopaminergic neurons and human iPSC-derived neuronal models will help establish the physiological and clinical relevance of this mechanism.

Together, our findings support a stepwise model in which mitochondrial stress promotes the initial accumulation of DJ-1 at mitochondria, followed by VDAC1-dependent enrichment within a protected submitochondrial compartment consistent with the matrix. In this model, the initial recruitment of DJ-1 may be driven by stress-associated changes in mitochondrial metabolism or redox state, whereas VDAC1 appears to facilitate the subsequent matrix-associated localization of DJ-1. Although the function of this matrix-associated pool remains to be fully defined, previous studies have implicated DJ-1 in Complex I activity or assembly ^31,33^ and mitochondrial ATP production ^9^. Importantly, inhibition of ATP synthase was sufficient to promote mitochondrial DJ-1 accumulation in the absence of an exogenous oxidative insult, suggesting that bioenergetic dysfunction itself is a key upstream trigger for DJ-1 redistribution. Furthermore, the ability of matrix-targeted DJ-1 to improve mitochondrial respiration, mitochondrial morphology and neuronal survival despite failing to restore spare respiratory capacity, supports a model in which matrix associated DJ-1 contributes substantially–but not exclusively–to mitochondrial protection during stress.

Overall, our findings identify a previously unrecognized VDAC1– DJ-1 axis that links mitochondrial stress sensing to compartment-specific DJ-1 localization and mitochondrial resilience. By demonstrating that VDAC1 supports stress-associated localization of DJ-1 to a protected mitochondrial compartment consistent with the matrix, and that restoring this pool improves mitochondrial respiration, morphology, and neuronal survival, our study provides new mechanistic insight into how compartment-specific DJ-1 localization preserves mitochondrial integrity during cellular stress.

### Limitations of the Study

Submitochondrial localization of DJ-1 was assessed using protease protection and biochemical fraction rather than live-cell imaging, an approach that reflects the distribution of DJ-1 across a population of mitochondria but does not resolve the dynamics of DJ-1 trafficking between compartments over time. In addition, because matrix targeting of DJ-1 in our rescue experiments was achieved using an engineered localization sequence, these experiments establish the functional importance of matrix-associated DJ-1 but do not establish the molecular mechanism by which endogenous DJ-1 is imported into the mitochondrial matrix or how VDAC1 regulates this process.

## Resource Availability

### Lead Contact

Requests for further information and resources should be directed to and will be fulfilled by the lead contact, Alvin Joselin.

## Materials Availability

This study generated new AAV constructs (AAV2/9-CBA-OMM-DJ1-myc and AAV2/9-CBA-MT-DJ1-myc). These reagents are available from the lead contact upon request.

## Data and Code Availability

All data reported in this study are included in the main text or supplemental information.

This study did not generate any new code.

Any additional information required to reanalyze the data reported in this paper is available from the lead contact upon request.

## Acknowledgements

This work was supported by grants from the Canadian Institutes of Health Research (grant numbers FRN #15123 and 184102) to D.S.P. S.J.H. was supported by a graduate award from Parkinson Society Canada and a doctoral award from the Canadian Institutes of Health Research (CIHR). A.J. acknowledges support from the Hotchkiss Brain Institute Investigator Research Award and the CaPRI Award, which support his Parkinson’s disease research program. We thank Dr. Frank Visser and the Molecular Core Facility of the Hotchkiss Brain Institute, University of Calgary, for the production of AVV particles used in this study. We thank Dr. William J. Craigen (Baylor College of Medicine) for providing VDAC1 MEFs and transgenic mice.

## Author Contributions

JT, SJH, DSI, and UA designed and performed experiments. DF performed the DJ-1 interactor screen. SMC and GK provided technical support. MB provided DJ-1 constructs. RSS, ZK, and NK contributed to experimental design and data interpretation. JT, SJH, DSI, and AJ analyzed and interpreted the data. JT, DSI, SJH, and AJ wrote the original manuscript, and all authors contributed to manuscript review and revision. AJ and DSP conceived and supervised the study, secured funding, and provided project administration. All authors read and approved the final manuscript.

## STAR Methods

### Experimental Model and Study Participant Details

#### Animals

The generation and genotype of DJ-1 and VDAC1 mice have been described previously ^6,23,3^ ^4^. All animal procedures were approved by the University of Ottawa Animal Care Committee and University of Calgary Animal Care Committee. Animals were maintained in accordance with the guidelines of the Canadian Council on Animal Care and institutional animal care protocols.

#### Cell culture

Mouse embryonic fibroblasts (MEFs) were derived from E14.5-15.5 transgenic mice of either sex on a C57BL/6 background and immortalized as previously described ^35^. VDAC1 WT and KO MEFs were generously provided by Dr. William Craigen and were immortalized by spontaneous selection of cells that persisted beyond typical senescence. All MEFs were maintained in Dulbecco’s Modified Eagle Medium (DMEM; Gibco, 11995-065) supplemented with 10% fetal bovine serum (FBS; Gibco, 12484-026) and antibiotic-antimycotic solution (Thermo Scientific, 15240062). Cultures were maintained at 37°C in a humidified incubator with 5% CO_2_.

Primary cortical neurons were isolated from E14.5-15.5 VDAC1 WT or KO mouse embryos of either sex and maintained in Neurobasal medium (Gibco, 21103049) supplemented with B27 containing antioxidants (Gibco, 17504-044), N2-supplement (Gibco, A1370701), 0.5 mM L-glutamine (Gibco, 25030-081), and penicillin-streptomycin (Gibco, 15140-122) as previously described^6^. Neurons were plated on 100 μg/mL poly-D-lysine-coated plates (Sigma-Aldrich, P0899) at the following densities: 5 x 10^6^ cells per 10-cm plate for mitochondrial fractionation; 1 x 10^5^ cells per well of a 24-well plate for survival assays and mitochondrial length analysis; 2 x 10^5^ per well of a 24-well plate for GFP-complementation assays; and 2 x 10^6^ per well of a 6-well plate for ATP assays.

### Method Details

#### AAV constructs

DJ-1 constructs targeted to the outer mitochondrial membrane (OMM-DJ-1) or mitochondrial matrix (MT-DJ1) were based on constructs described in Calì et al,, 2015^9^, provided by Dr. Marisa Brini (University of Padova). AAV2/3-CBA-GFP (2.35 × 10¹² GC/mL), AAV2/3-CBA-OMM-DJ1-GFP (6.63 × 10¹² GC/mL), and AAV2/3-CBA-MT-DJ1-GFP (6.63 × 10¹² GC/mL) were used as indicated. A second set of constructs was produced as AAV2/9 particles by the Molecular Core Facility of the Hotchkiss Brain Institute (University of Calgary), including AAV2/9-CBA-EGFP (4.52 × 10¹² GC/mL), AAV2/9-CBA-OMM-DJ1-myc (1.34 × 10¹³ GC/mL), and AAV2/9-CBA-MT-DJ1-myc (6.35 × 10¹² GC/mL), used as indicated. Neurons were transduced at the time of plating at an MOI of 500.

#### Mitochondrial enriched fraction and trypsin digestion

VDAC1 WT and KO MEFs were treated up to 3h with 100μM H_2_O_2_ and then subjected to fractionation as previously described ^7^. Similarly, VDAC1 KO cortical neurons from E14.5-15.5 mouse embryos were cultured for 5d in the presence of antioxidants. Cortical neurons received new media without antioxidants at DIV5. At DIV6, cortical neurons were treated with either 10 μM MPP^+^ (Sigma-Aldrich, D048) up to 24 h or 10 μM oligomycin (Sigma-Aldrich, O4876) up to 1 h and subjected to fractionation. Because the primary objective of these experiments was to compare WT and VDAC1 KO samples at each predefined treatment time point, genotype comparisons were performed using two-tailed unpaired Student’s t-tests. For antioxidant experiments, neurons were pre-treated with 2 mM N-acetylcysteine (NAC; Sigma-Aldrich, A7250) for 2 h prior to stress. For trypsin digestion, VDAC1 WT and KO MEFs treated with 100 µM H_2_O_2_ for 3 h were subjected to a modified subcellular fractionation. Specifically, isolation buffer consisted of 200 mM mannitol, 70 mM sucrose, 10 mM HEPES, and 1 mM EGTA with pH 7.5 and no protease inhibitors were added. The mitochondrial-enriched fractions were digested with 25 μg/mL trypsin (Sigma-Aldrich, T8253) for up to 120 min. To stop the digestion, 50 μg/mL of AEBSF (Thermo Scientific, 78431) was added to the reaction. For membrane permeabilization controls, mitochondrial fractions were incubated with 1% Triton X-100 prior to trypsin digestion.

#### Co-immunoprecipitation (Co-IP)

Co-IP was performed in human embryonic kidney (HEK) 293 cell line. GST-DJ1 or GST alone was transiently transfected using Lipofectamine 2000 (Gibco, 11668-019) and cells were lysed for 24-36 h in lysis buffer (50 mM Tris HCl pH 7.5, 100 mM NaCl, 1 mM EDTA, 1 mM DTT, 0.2% NP-40 and protease inhibitor). Lysate was incubated with 40 μL of glutathione Sepharose (GE Healthcare, 17-0756-05) for 4 h. Samples were washed 3 times with lysis buffer. Bound proteins were eluted by boiling samples in 2X SDS-loading buffer and analyzed by Western blot. Endogenous immunoprecipitation of DJ-1 was performed in mitochondrial-enriched fractions from either VDAC1 WT and VDAC1 KO MEFs, or DJ-1 WT and DJ-1 KO MEFs. Mitochondrial-enriched fractions were resuspended in isolation buffer and pre-cleared with a mixture of Protein A agarose beads (Sigma-Aldrich, P9424) and Protein G agarose beads (GE Healthcare, 20421) for 6h. Cleared lysate was incubated with 5 μg of DJ-1 antibody (Abcam, ab18257) or normal mouse IgG (Santa Cruz, SC - 2025) overnight. Samples were washed 5 times with isolation buffer and eluted by boiling in 2X SDS-loading buffer. All Co-IP experiments were independently repeated at least three times for HEK293 and MEF samples.

#### Western blot

Protein quantification was performed using Bradford (Bio-Rad, 5000006) method. Samples were electrophoresed on 12% sodium dodecyl sulfate polyacrylamide gels and transferred to polyvinylidene fluoride (PVDF) membranes (Millipore, IPFL00005). Membranes were probed with the respective primary antibodies followed by horseradish peroxidase– conjugated secondary antibodies and developed with the Immobilon Western Chemiluminescent HRP Substrate (Millipore, WBKLS0500). Densitometry analysis was carried out using ImageJ (NIH). For mitochondrial-enriched samples 10 μg of each sample was analyzed. For mitochondrial digestion samples, equal volumes of sample were assessed. Primary antibodies included: for the OMM - Tom20 (Abcam, ab78547, rabbit, 1:2000); for the IMM - NDUFA9 (Thermo Scientific, 459100, mouse, 1:2000); for the matrix and loading control - mtHSP70 (Abcam, ab2787, mouse, 1:2000); VDAC1 (Santa Cruz, sc390996, mouse, 1:2000); DJ-1 (Abcam, ab18257, rabbit, 1:20,000); GST (Santa Cruz, sc33613, rabbit, 1:2000). Secondary antibodies included: HRP-conjugated goat anti-mouse IgG (H+L) (Bio-Rad, 1706516, 1:10000); HRP-conjugated goat anti-rabbit IgG (H+L) (Bio-Rad, 1706515, 1:10000); TrueBlot anti-mouse IgG HRP (Rockland, 18-8817-31, 1:1000); TrueBlot anti-rabbit IgG HRP (Rockland, 18-8816-33, 1:1000).

#### Immunofluorescence

Following treatment, neurons were fixed in 4% formalin in growth media at room temperature for 30 min. The cells were permeabilized and blocked simultaneously using 0.1% Triton X-100 and 5% normal goat serum for 1 h at room temperature. Incubation with the respective primary antibodies (DJ-1, Abcam, 1:500; Tom20, Abcam, 1:500; COXI, Abcam, ab14705, 1:300) was performed overnight at 4°C. Samples were subsequently washed in 10% normal goat serum and stained with the corresponding secondary antibodies (1:500). Following 3 washes in phosphate-buffered saline (PBS), samples were stained with Hoechst and analyzed by confocal microscopy.

#### Measurement of Mitochondrial length

Mitochondrial length was assessed using ImageJ (NIH) as previously ^5^. Briefly, length of mitochondria stained for Tom20 were measured in VDAC1 WT and KO cortical neurons transduced with AAV2/3-CBA-GFP, AAV2/3-CBA-OMM-DJ1-GFP or AAV2/3-CBA-MT-DJ1-GFP with control and 10 μM MPP^+^ treatment. Lengths were either averaged or binned accordingly. A minimum of 500 mitochondria were counted per sample size (n). Cell survival assay was carried out as described previously ^35^. Briefly, cortical neurons were fixed in 4% formalin in growth media at room temperature for 30 min after 48 h of 10 μM MPP^+^ treatment. Subsequently they were stained with Hoechst and the number of apoptotic nuclei was analyzed by fluorescence microscopy.

#### ATP assay

The CellTiter Glo Luminescent (Promega, G7570) assay was performed for a measurement of ATP level in the cell as described previously ^9^. Briefly, following 10 μM MPP^+^ treatment, cortical neurons were harvested by gentle trypsinization using TrypLE reagent (Invitrogen, 12563011). Each sample was divided into 6 wells of an opaque 96-well plate, of which 3 wells were incubated with 10 μM oligomycin for 1 h to inhibit mitochondrial-produced ATP. An aliquot of each sample was used to determine cell number by a hemocytometer. Luminescence was recorded using a plate reader. A range of standard ATP concentrations was used to determine ATP concentrations in samples. Data is presented as μM of ATP corrected for cell number.

#### Reactive Oxygen Species (ROS) assay

Cortical neurons were seeded onto poly-D-lysine–coated 24-well plates at 100,000 cells per well. At DIV6, neurons were treated with 10 μM MPP⁺ in antioxidant-free neurobasal medium for 1 or 3 h. After treatment, cells were incubated with 10 μM CM-H₂DCFDA (Abcam, ab113851) for 30 min at 37°C, washed, and imaged by wide-field fluorescence microscopy. For quantification, mean fluorescence intensity was measured in 4–6 randomly chosen fields per condition using ImageJ. Each condition was repeated in 3 independent biological replicates.

#### GFP complementation system

Bi-fluorescent complementation system was used for submitochondrial localization of DJ-1^9^ . Briefly, cortical neurons at DIV 4 were transfected with a small part of GFP fused with DJ-1 (WT-DJ1-S11) and a complementary part of GFP targeted to either the OMM (OMM-GFP1-10) or matrix (MT-GFP-10) using Lipofectamine 3000 (Thermo Scientific L3000008). 24 h later, media was changed to neurobasal media without antioxidant and neurons were treated with 10 μM MPP^+^ for 3 h and fixed. Samples were stained with COXI to label mitochondria. The relative level of GFP to mitochondrial fluorescence was measured for localization of DJ-1.

#### Measurement of mitochondrial oxygen consumption rate (OCR)

The Seahorse XF24 Extracellular Flux Analyzer (Seahorse Biosciences) was utilized to evaluate oxygen consumption in cortical neurons as described previously ^36^. The cortical neurons were seeded onto poly-d-lysine-coated 24-well seahorse plates at a density of 1.5 × 10^5 cells/ well in 500 μL neurobasal media supplemented with 1% B27, glutamine, and penicillin-streptomycin. For each Seahorse experiment, cortical neurons were derived from individual E14.5–15.5 embryos (WT or KO), and each biological replicate corresponds to one embryo-derived culture. Neurons were plated at equal densities on Seahorse XF-24 plates and analyzed after 9–10 days in vitro. Following 9–10 days of culture, the medium on the neurons was replaced with bicarbonate-free assay buffer (120 mM NaCl, 3.5 mM KCl, 0.4 mM KH₂PO₄, 5 mM NaHCO₃, 20 mM HEPES, 1.2 mM Na₂SO₄, 1.2 mM CaCl₂, and 1–2 mM MgCl₂, pH 7.4) supplemented with 5 mM glucose, 4 mM glutamine, 1 mM pyruvate, as previously described ^37^, and was incubated for 30 min in a CO_2_-free incubator prior to being loaded into the XF Analyzer. The basal respiration was measured first, followed by the sequential treatment of oligomycin (500 ng/mL) to assess ATP-linked respiration, FCCP (2 μM) to determine maximal respiration capacity, and Rotenone (0.5 µM)/ antimycin A (1 μM) to measure non-mitochondrial OCR. Each measurement was taken over a 2-min interval, followed by 2 min of mixing and 2 min of incubation. Four measurements were taken for basal OCR, and three measurements were taken after oligomycin, FCCP, and rotenone/ antimycin A treatment. The data was compiled by the XF software, normalized to protein levels per well, and final data prepared using Microsoft Excel. The experiment was repeated at least three times independently using different litters, with consistent outcomes across replicates.

For the rescue experiments, oxygen consumption was measured using a Seahorse XF96 Extracellular Flux Analyzer (Agilent/Seahorse Biosciences). VDAC1 KO cortical neurons transduced with AAV2/9-CBA-EGFP, AAV2/9-CBA-OMM-DJ1-myc or AAV2/9-CBA-MT-DJ1-myc were seeded onto poly-D-lysine and laminin (5 μM/mL; Sigma L2020)-coated XF96 plates (Agilent, 103793-100) at a density of 70,000 cells/ well in 200 μ L neurobasal media supplemented with 1% B27, glutamine, and penicillin-streptomycin. Neurons were analyzed after 8 days in vitro, in bicarbonate-free assay buffer, following sequential injection of oligomycin (1 μM), FCCP (2 μM), and rotenone/antimycin A (0.5 μM) (Agilent, 103015-100). Data were normalized normalized to protein levels per well. This experiment was performed independently twice, with n = 4 per group in each.

#### Quantification and Statistical Analysis

Statistical analyses were performed using GraphPad Prism version 8.0. For comparisons between two groups, statistical significance was determined using two-tailed Student’s t test. For comparisons among multiple groups, one-way or two-way analysis of variance (ANOVA) was used, followed by Tukey’s post hoc test where appropriate. Data are presented as mean ± standard error of the mean (SEM). No blinding or randomization was performed. Sample sizes were not pre-determined by a power calculation but were based on those commonly used for similar experiments in the field. Statistical details of each experiment, including the value of n and the statistical test used are reported in the corresponding figure legends.

**Supplementary Figure 1. DJ-1 associates with VDAC1 in embryonic mouse brain.**

Endogenous co-immunoprecipitation of DJ-1 and VDAC1 from embryonic brain lysate. Lysate was immunoprecipitated with anti-VDAC1 antibody or control IgG and analyzed by Western blotting for DJ-1 and VDAC1. Input lysate is shown as a positive control. The asterisk indicates the VDAC1-specific band. A non-specific IgG light-chain band is visible below the VDAC1-specific band. Representative immunoblots from n = 1 experiment.

**Supplementary Figure 2. Membrane permeabilization abolishes protease protection of mitochondrial DJ-1.**

Mitochondrial-enriched fractions from VDAC1 WT MEFs were treated with trypsin for the indicated durations in the absence (A) or presence (B) of 1% Triton X-100 before digestion. Samples were immunoblotted for DJ-1 and mitochondrial compartment markers Tom20 (outer mitochondrial membrane), NDUFA9 (inner mitochondrial membrane), and mtHSP70 (mitochondrial matrix). Representative immunoblots from n = 3 independent experiments.

