## Supplementary figures and images for "VDAC1 regulates stress-associated matrix localization of DJ-1 to support mitochondrial homeostasis and neuronal survival"

### Supplementary Figure 1

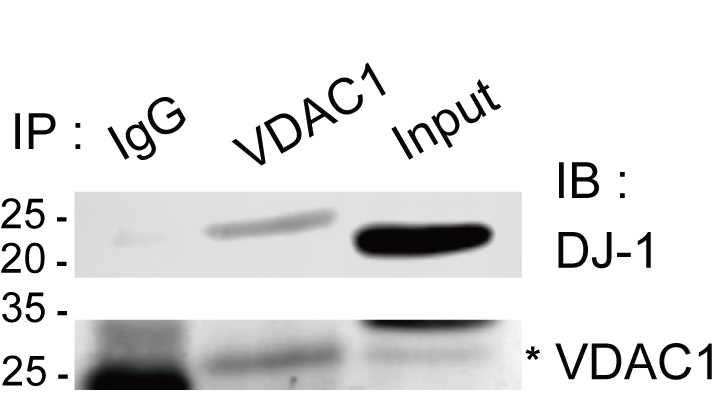

### Supplementary Figure 2

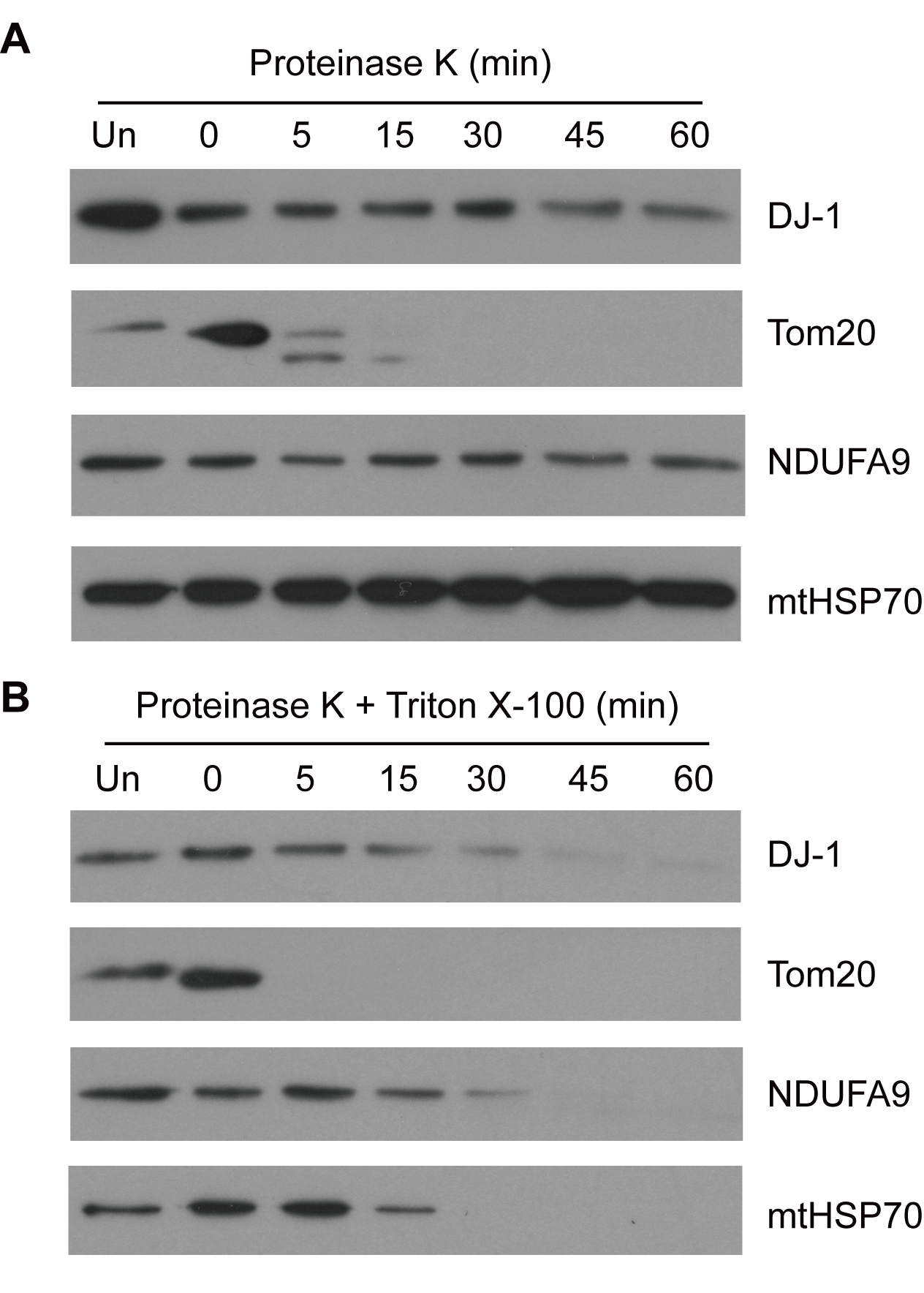
